# Self-organization of human dorsal-ventral forebrain structures by light induced SHH

**DOI:** 10.1101/2021.08.18.456849

**Authors:** Riccardo De Santis, Fred Etoc, A. Edwin Rosado-Olivieri, Ali H. Brivanlou

## Abstract

Organizing centers secrete morphogens that specify the emergence of germ layers and the establishment of the body’s axes during embryogenesis. While traditional experimental embryology tools have been instrumental in dissecting the molecular aspects of organizers in model systems, they are impractical in human *in-vitro* model systems to quantitatively dissect the relationships between signaling and fate along embryonic coordinates. To systematically study human embryonic organizer centers, we devised a collection of optogenetic ePiggyBac vectors to express a photoactivatable Cre-loxP recombinase, that allows the systematic induction of organizer structures by shining blue-light on hESCs. We used a light stimulus to geometrically confine SHH expression in neuralizing hESCs. This led to the self-organization of mediolateral neural patterns from the organizer. scRNA-seq analysis established that these structures represent the dorsal-ventral forebrain, at the end of the first month of development. Here, we show that morphogen light-stimulation is a scalable tool that induces self-organizing centers.

## Introduction

During embryogenesis, the central nervous system (CNS) is derived from an epithelial sheet of cells, the embryonic neural plate, that is induced in a polarized non-cell autonomous manner by a small group of cells called the Spemann organizer (Spemann and Mangold, 1924). Neural induction activity of the organizer occurs by a default mechanism that is exerted through the secretion of soluble inhibitors that block both branches (SMAD1/5/8 and SMAD2/3) of the TGFb signaling pathway (Harland and Gerhart, 1997; Sasai et al., 1994; Brivanlou et al., 1994; Ozair et al., 2013; De Santis and Brivanlou, 2020). Dual SMADs inhibition directly converts pluripotent embryonic stem cells to the anterior neural tissue of the dorso-anterior forebrain (Chambers et al., 2009). Downstream of these primary neural inducing signals, highly localized and dynamic organizing centers provide multiple morphogen sources that pattern the central nervous system (Arnold and Robertson, 2009; Borello and Pierani, 2010; Kiecker and Lumsden, 2012). The embryonic neural tissue undergoes antero-posterior (A-P) and medio-lateral (M-L) patterning during neural plate stages (Cox and Brivanlou 1995; Ozair et al., 2013; Nikolopoulou et al., 2017). As the neural tube closes, cells in the lateral part of the plate become dorsal (roof plate) and those in the midline become ventral (floor plate). Thus, M-L patterns are converted to dorso-ventral polarity (D-V).

The interplay between two signaling pathways, BMP4 and SHH, both acting as morphogens guide the establishment of the M-L and D-V polarity of the embryonic neural tissue. BMP4, is expressed in the lateral edge of the neural plate, and subsequently in the dorsal neural tube, while SHH, is expressed in the ventral midline of the neural plate, and subsequently in the floor plate of the neural tube (Lupo et al., 2006; Ozair et al., 2013). Classical experimental embryology approaches such as ectopic presentation of SHH ligand by grafting coated beads, embryonic explants, or morphogen-secreting cells in mouse, chick or human stem cells demonstrated that SHH activity is sufficient to induce ventral neural fates via its transcriptional effector Gli3 (Kuschel et al., 2003; Marklund et al., 2004; Lupo et al., 2006, Maroof et al., 2013; Placzek and Briscoe, 2018, Cederquist et al., 2019).

While these approaches have been instrumental in shaping our current understanding, they also suffer from technical shortcomings that have hindered a precise mapping of fate acquisition as a function of signaling dynamics in the context of early human development. For example, grafting experiments provide little control over the extent of the inductive field’s spatial limits, control of throughput, and non-specific effects due to wound healing. The development of new tools that systematically control these parameters will lead to better dissection of morphogens patterning and will constitute a major step forward in experimental embryology. Optogenetic tools have been recently used to control gene expression with spatiotemporal resolution, taking advantage of different strategies (Kennedy et al., 2010; Kawano et al., 2016; Nihongaki et al., 2017; Quejada et al., 2017; Shao et al., 2018; Morikawa et al., 2020; Rogers and Muller, 2020). This carries the potential of creating exogeneous embryonic organizer centers in model tissues for quantitative studying of embryonic induction and for creating *in-vitro* self-organized structures that present the axial organization, that is a key landmark of embryonic development. Light modulation of signaling pathways provides flexibility and high spatial resolution over the morphogenetic stimulus (Kim et al., 2014; Sako et a., 2016; Johnson et al., 2017; Rogers and Muller, 2020; Repina et al., 2020).

Here, we have devised a collection of optogenetic ePiggyBac vectors to conditionally express a photoactivatable Cre-loxP recombinase for creating spatially restricted organizing centers that break symmetry in self-organizing hESCs. This collection is an hESCs optimized blue-light inducible split-CRE system based on the Magnets-split-CRE (Kawano et al., 2016). This system allows precise spatio-temporal control over the expression of a morphogen under a blue-light input. This experimental set up provides a highly quantitative and simplified method based on blue-light stimulation that can be used to induce organizing centers in *in-vitro* cultures of hESCs. To establish proof of feasibility, we tested our tool for its ability to break symmetry in hESC-derived neural tissue to light-induce M-L polarity by inducing stripes of SHH expression as observed in the midline of the neural plate in embryo. Light induced and polarized expression of SHH during neural induction in absence of exogenous WNT modulation, led to the self-organization of a 2D *in-vitro* human dorsal-ventral forebrain structure, that include a ventral telencephalic-hypothalamic primordia. Polarization of morphogens using light provides a non-invasive approach to decipher the earliest events that underly symmetry breaking in the embryonic nervous system in stages of human development otherwise inaccessible for scrutiny.

## Results

### Engineering a collection of optogenetic ePiggyBac vectors to conditionally express a photoactivatable Cre-loxP recombinase in hESCs

In order to provide light modulation of gene expression to human developmental studies, we re-engineered our original transposon ePiggyBac vector (Lacoste et al., 2009, Rosa et al., 2014) to conditionally express a light-inducible Cre-recombinase enzyme that takes advantage of the Magnets dimerization system (Magnet-CRE) (Kawano et al., 2016) (Fig 1A). This allows for a stable integration of a blue-light dependent CRE enzyme in the genome. To avoid culturing cells in the dark, minimize leakage and to gain better control of light-sensitivity, we controlled the light-CRE enzyme using a Dox-inducible promoter and a second T2A peptide to improve the separation of its components (Fig 1A, left panel). We paired this vector with a receiver ePiggyBac that carries two sequential ORFs (Red and Green modules) to be regulated by LoxP recombination in a mutually exclusive manner (Fig 1A, right panel). Both vectors were stably transfected into our female XX hESC line, RUES2 (NIHhESC-09-0013) and stimulated with DOX and blue-light through a light-blocking photomask (Fig 1B, Suppl. Fig 1A-B-C). Dox- and Blue-light induced hESCs showed robust expression of the Green module in patterns imposed by the shape of the photomask (Fig 1B-C, Suppl. Fig1B-C). The Green module expressing cells showed a highly reproducible pattern of expression that tightly correlate with the photomask that slightly increase over time, probably due to cell proliferation (Suppl. Fig 1D). The efficiency of light conversion by single cell fluorescence measurement shows that 24 hours of blue-light stimulation converts the 78.3 % of the total cells, while controls kept in the dark, or in absence of Dox, show less than 1% of Green positive cells (Fig 1D). The activation of the green module depends on the duration of blue-light stimulation. Pulsed blue-light for 600 cycles (24 hours) is the most efficient treatment without inducing cell death, as measured by CASP3 activation (Fig 1D, Suppl. Fig 1E). The blue light-dependent induction of the Green module was also validated by measuring RNA levels by qRT-PCR in whole-illuminated samples (Fig 1E). LoxP genomic recombination upon Dox and light treatment is shown by amplicon Sanger sequencing of the selected genomic region (Fig 1F). The light induced pattern of gene expression is consistently and stably maintained over six days in the absence of continuous light stimulation (Suppl. Fig 1F). Light induced gene expression modulation was not confined to a single hESC line as another of our hESC line (RUES1, genetic background, male XY, NIHhESC-09-0012) responded in the same manner (Suppl. Fig1G). Collectively, these experiments demonstrate that pairing a light inducible CRE enzyme with a stable and drug inducible ePiggyBac vector allows for a rapid and efficient spatiotemporal control of exogenous gene expression in hESCs.

**Figure 1:**
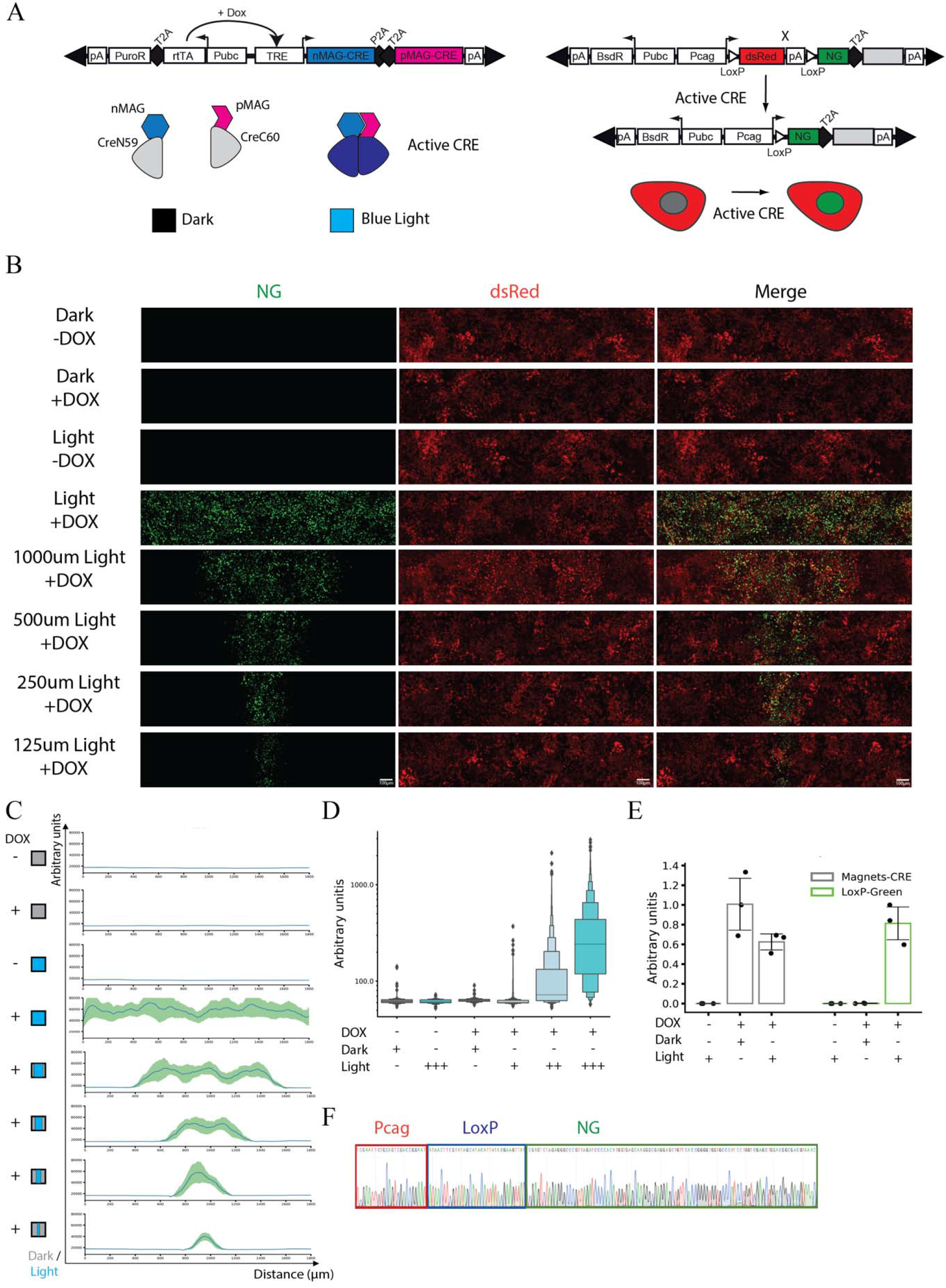
Light-induced gene expression programs in hESCs. **(A)** Schematization of the collection of optogenetic ePiggyBac vectors to conditionally express a photoactivatable Cre-loxP recombinase vector. Left panel. Puromycin selectable and Dox inducible piggyBac transposon (Lacoste et al., 2009; Rosa et al., 2014) carrying the CRE-MAGNETs system (Kawano et al., 2016). Dox treatment induces the CRE-MAGNETs system which reconstitutes a fully active CRE in the presence of blue light. Right panel. Blasticidin selectable LoxP exchangeable dual colors piggyBac transposon. The first ORF (Red module) is constitutive expresses while the second ORF (Green module) depends on CRE-LoxP recombination. Grey Box is the possible cargo protein taking advantage of the T2A peptide. **(B)** Photomask induced light patterns showing the expression of the Red module (dsRed), Green module (NG) and merge composite channel. Dark control and spatially localized activation in presence and absence of DOX with different spatial features are shown (1000 μm, 500 μm, 250 μm, 125 μm). Scale bar: 100um. **(C)** Cumulative fluorescence intensity analysis (line profile) over the x-axis. x-axis displays the linear distance in μm. y-axis reports the cumulative fluorescent intensity profile in arbitrary units. Line profile shows the average (line) and standard deviation (area) for the Green fluorescent channel. Quantification after 600 cycles (24 hours) of pulsed Blue light (n=3). **(D)** Single-cell fluorescent intensity quantification. Conditions: presence or absence of Dox with concomitant Dark or Light stimulation. We titrated the time of light stimulation using different pulsed light conditions. 1 cycle of pulsed light is equal to 20sec Light-ON and 120sec Light-OFF. The light stimulation intervals are: 1 cycle, 25 cycles (1 hour), and 600 cycles (24 hours) (n=5). **(E)** qRT-PCR showing mRNA induction of the MAGNETs system and light-induced expression of the Green module (n=3). **(F)** Sanger sequencing of the genomic region flanking the LoxP site showing the recombination of the Red module and the generation of the Green module (Red box: Pcag-promoter, Blue box: LoxP, Green box: NG).

### Generation of dorso-ventral polarity in hESC-derived neural tissue

In order to study the M-L and D-V aspects of neural patterning in human models, we tested the ability of light stimulation to generate a localized SHH organizing center. We devised a LoxP inducible Green module to be co-expressed with SHH at the mRNA level, while producing two separate proteins using a T2A peptide. This setup can be regulated by blue-light stimuli through LoxP recombination (Fig 2A). hESCs (RUES2) were differentiated using dual SMADs inhibition (SB431542 and LDN193189) to induce an anterior forebrain fate. The application of DOX at day 0 confers light sensitivity for the first two days of differentiation (Fig 2A). Neural induction using dual-SMADs inhibition has been shown to generate progenitors representative of the embryonic neuroepithelium (Chambers et al., 2009; Ozair et al., 2019; Haremaki et al., 2020). At this stage, a set of defined markers can be used to decipher patterning at various distances from the SHH source: PAX6 marks the most dorsal population, FOXA2 the floor plate and NKX2.1 the ventral neural progenitors in the basal telencephalon and hypothalamic primordia at E12.5 (Suppl. Fig 2A) (Fasano et al., 2010, Maroof et al., 2013).

**Figure 2:**
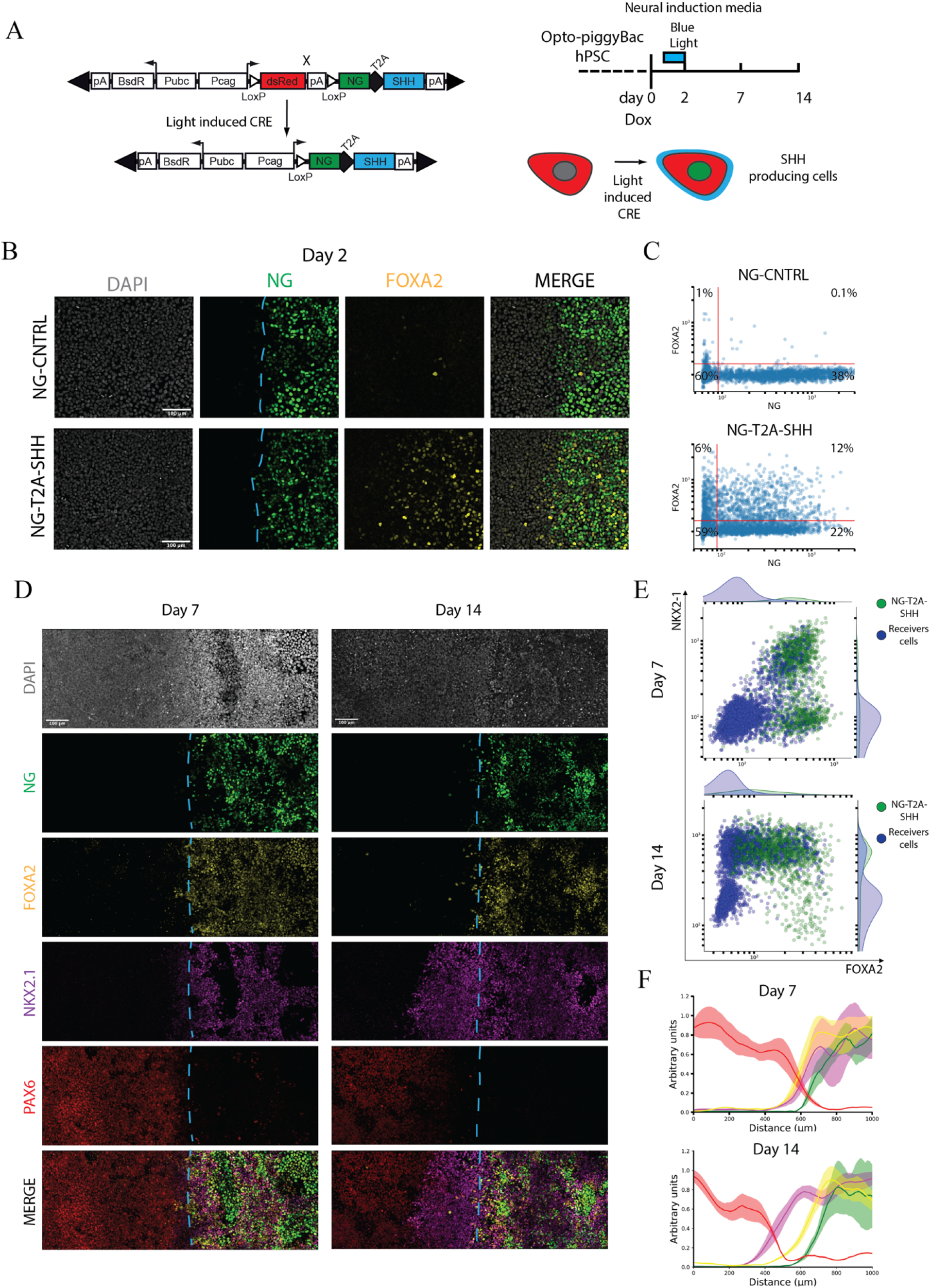
Medio-lateral neural patterning by light-induced SHH. **(A)** Left panel. Schematization of Green module T2A human SHH. This setup allows conditional expression of SHH and fluorescent visualization of the light-induced cells. Right panel. Schematization of Blue-light stimulation and neural induction reporting analyzed time points. Neural induction is induced by inhibiting TGFβ signaling by dual-SMADs inhibition that is maintained during the entire course of differentiation (14 days). Dox treatment starts at day “0” and it is wash-out at day “2”. **(B)** FOXA2 immunostaining in light induced SHH cells and NG-CNTRL during differentiation at day 2. DAPI-Grey, NG-Green, FOXA2-Yellow, Merge-Composite. Dashed cyan line indicate the border of SHH producing cells. Scale bar: 100μm. **(C)** Day 2 single cell fluorescence quantification displayed as a scatterplot, reporting FOXA2 and NeonGreen (NG) intensity for NG-CNTRL and NG-T2A-SHH. **(D)** Immunostaining time-course analysis of the dorsal and ventral fates, revealing NG-Green (Organizer), FOXA2-Yellow (Ventral floor plate), NKX2-1-Magenta (Ventral neural progenitors), PAX6-Red (Dorsal neural progenitors), Merge-Composite at day 7 and day 14 during neural differentiation in response to ligh patterned SHH. Dashed cyan line indicate the border of SHH producing cells. Scale bar: 100μm. **(E)** Single cell fluorescence quantification of NG-T2A-SHH induced cells displayed as combined scatterplot and density histogram at day 7 (top) and day 14 (bottom). x-axis report FOXA2 while the y-axis the NKX2-1 fluorescence intensity profile. Each dot represents an individual cell that has been color coded according to its green module status. The green dots represent the NG-T2A-SHH positive cells, while the blue dots are NG-T2A-SHH negative cells. (**F)** Cumulative fluorescence intensity analysis (line profile) over the x-axis. x-axis displays the linear distance in μm. y-axis reports the cumulative fluorescent intensity profile in arbitrary units for each channel. Line profile shows the average (line) and standard deviation (area) for each channel. The line profile is color-coded as the immunofluorescent channels, NG-Green, FOXA2-Yellow, NKX2-1-Magenta, PAX6-Red. Top panel. Day 7 quantification (n=4). Bottom panel. Day 14 quantification (n=4).

Blue light was shone on day 1 during neural induction through a 1 mm rectangular mask for 24 hours to produce SHH and a green fluorescent protein (NG-T2A-SHH) or a control fluorescent protein (NG-CNTRL) in a spatially restricted domain. Light stimulation induced the expression of the Green module in both NG-CNTRL and NG-T2A-SHH lines (Fig 2B). At day 2, FOXA2 positive cells were specifically induced by the NG-T2A-SHH but not in the NG-CNTRL line (Fig 2B-C, Suppl. Fig 2B). FOXA2 positive cells were detected in cells secreting SHH as well as in cells just next to the SHH secreting domain, providing functional evidence of autocrine as well as paracrine SHH activity (Fig 2B-C, Suppl. Fig 2B-C). Examination of the light-induced cells after 7 and 14 days display FOXA2^+^, NKX2-1^+^ and PAX6^+^ cells that are organized in discrete domains while NG-CNTRL cells acquire PAX6^+^ default neural fate, in absence of ventral cell types (Fig 2D-E-F; Suppl. Fig 3A-B-C). The expression NG-T2A-SHH is stably maintained in absence of light over the course of the differentiation (Suppl. Fig 3D-E). SHH light induction unveiled M-L self-organization of the neural populations under the influence of an organizing center. Interestingly, at day 7, a population of cells co-expressing FOXA-2 and NKX2-1 was detected within and near by the light-induced organizer (Fig 2D-E-F, Suppl. Fig 3A). These FOXA2^+^/ NKX2-1^+^ double-positive cells have been suggested to be human specific, as have not been detected in the mouse, while they are present in the ventral forebrain in human fetal samples at PCW5.5 (Maroof et al., 2013). PAX6, NKX2-1 and FOXA2 domains gradually segregate over time inside and outside the SHH induced domain, with PAX6^+^ cells localized the farthest from the SHH source (Fig 2D-F). Single cell quantification shows that at day 7, the organizer induces population of cells double positive for NKX2-1^+^/FOXA2^+^, both cell autonomously and non-cell autonomously (Fig 2E, upper panel). A fraction of light converted cells, express high levels of FOXA2 but not NKX2-1 (Fig 2D-E, upper panel). At day 14, the ventral cellular populations induced by SHH differentiate into a NKX2-1^+^/FOXA2^-^ population, that is located both laterally and inside the light induced SHH organizer (Fig 2D-E, lower panel). Also, there is a population of cells NG-T2A-SHH^+^/FOXA2^+^ /NKX2-1^-^ (Fig 2E-F, Suppl. Fig 4A). NKX2-1 domain juxtaposed to the SHH organizer is induced independently from the size of the SHH domain (Suppl. Fig 4B). The RNA expression of the NG, the exogenous SHH and its downstream target GLI1 correlate with the expression of the NG-T2A-SHH module, validating our co-expression strategy (Suppl. Fig 4C). Therefore, our analysis revealed a proximal distal pattern of ventral cell fates from the SHH organizer during neural induction. Spatiotemporal control of SHH induces ventral neural fates that are organized in a 2D space *in-vitro*, resembling M-L and D-V neural populations (Suppl. Fig 2D). Moreover, it validates the functionality of our optogenetic tool for its ability to induce and self-organize discrete fates in hESCs with a simple blue light stimulation.

**Figure 3:**
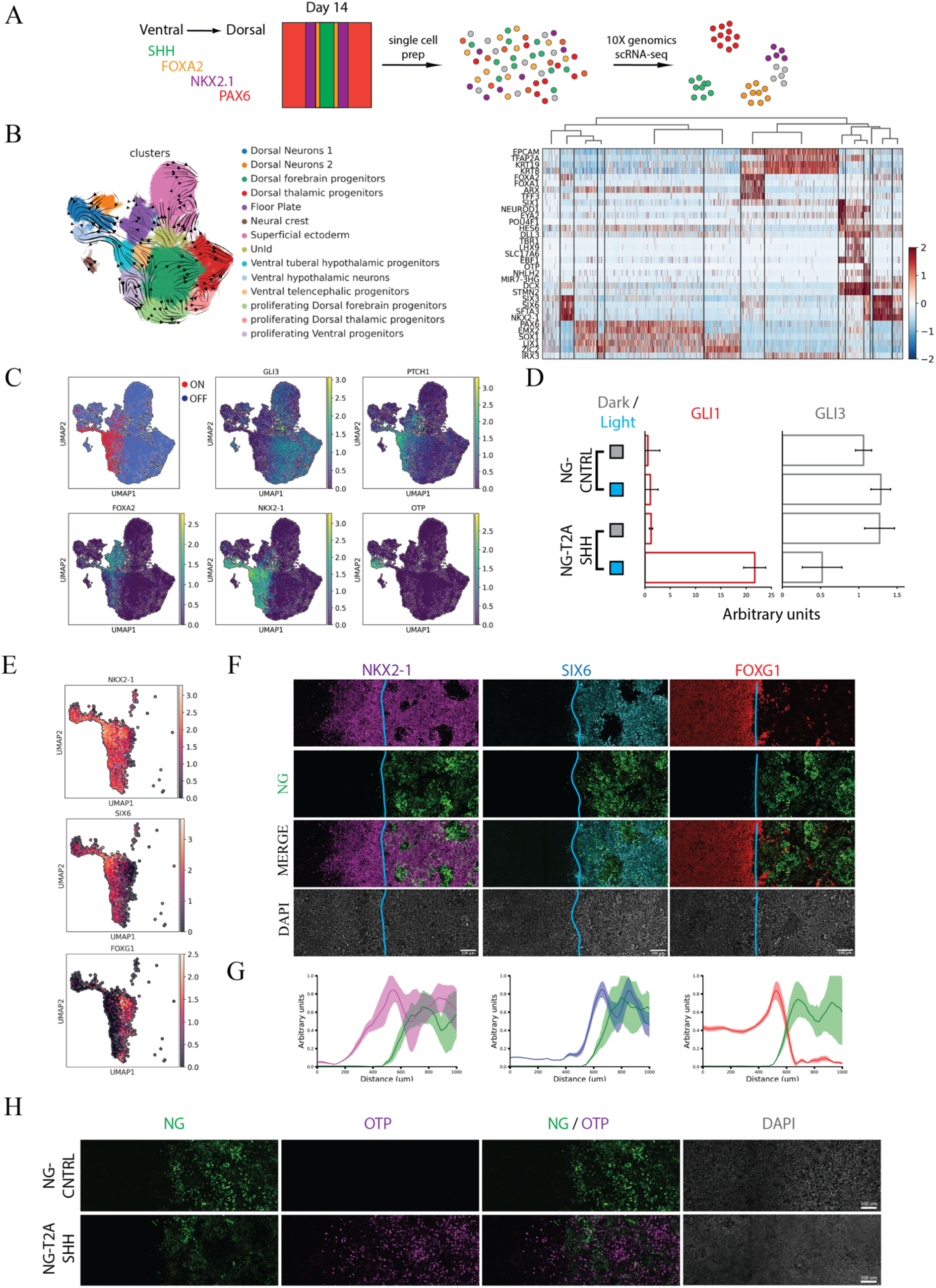
Spatial self-organization of telencephalic and hypothalamic fates upon light-induced SHH. **(A)** Schematization of the scRNA-seq profiling strategy. **(B)** Left panel: UMAP plot labeled with the identified cell identities and RNA velocity vectors: (i) FOXA2^+^/ARX Floor Plate; (ii) NKX2-1^+^/RAX^+^/SIX6^+^ Ventral tuberal hypothalamic progenitors; (iii) NKX2-1^+^/FOXG1^+^ Ventral telencephalic progenitors, (iv) NKX2-1^+^/NHLH2^+^/OTP^+^ Ventral hypothalamic neurons; (v) TFAP2A^+^/KRT19^+^ Superficial ectoderm; (vi) SOX10^+^/PLP1^+^ Neural Crest, (vii) PAX6^+^/EMX2^+^/OTX2^+^ Dorsal forebrain progenitors; (viii) IRX3^+^/ OLIG3^+^ Dorsal thalamic progenitors; (ix) TBR1^+^/LHX1^+^ Dorsal Neurons_1; (x) HES6^+^/DLL3^+^ Dorsal Neurons_2; dorsal and ventral proliferating progenitors, (xi) Dorsal, (xii) Dorsal thalamic, (xiii) Ventral and (xiv) an UnIdentified population (UnId). Right panel: z-score scaled heatmap of marker genes used for cell identities classification. **(C)** UMAP plot displaying SHH responsive and unresponsive status, and GLI3 and PTCH1 expression level. SHH responsive and unresponsive status have been computed imposing a threshold based on the normalized distribution of GLI3, GAS1 and PTCH1 counts, where GLI3-GAS1^low^ and PTCH1^high^ represent SHH responsive while GLI3-GAS1^high^, PTCH1^low^ unresponsive. **(D)** qRT-PCR analysis of SHH responsive genes (GLI1 and GLI3) for NG-T2A-SHH and NG-CNTRL at day 14 of differentiation. The histogram displays the average and standard deviation of differentiated cells exposed to light stimulation or dark control (n=2). **(E)** UMAP plot showing a selected population of cell NKX2-1^+^. Expression of marker that distinguish hypothalamic and telencephalic populations (NKX2-1, SIX6 and FOXG1). **(F)** Immunostaining shows the spatial segregation of ventral population arising from a light-induced SHH source at day 14. (Green NG, Magenta NKX2.1, Cyan SIX6, Red FOXG1, Grey DAPI), Scale bar = 100μm. **(G)** Cumulative fluorescence intensity analysis (line profile) over the x-axis. x-axis displays the linear distance in μm. y-axis shows the cumulative fluorescent intensity profile in arbitrary units for each channel. Line profile shows the average (line) and standard deviation (area) for each channel. The line profile is color-coded as the immunofluorescent channels, NG-Green, NKX2-1-Magenta, SIX6-Cyan, FOXG1-Red at day 14 quantification (n=3). **(H)** Immunostaining shows OTP positive cells induced in proximity of the NG-T2A-SHH organizer but not in the NG-CNTRL. Light induced SHH drive the self-organization of both neural progenitors and neurons (Green NG, Magenta OTP, Grey DAPI), Scale bar = 100μm.

### Molecular characterization of light-induced ventral neural fate

In order to precisely and unbiasedly identify the cell types present in our light-induced, self-organizing neural tissue, we characterized their transcriptome using scRNA-seq from 2 independent biological replica (12097 and 6207 cells) (Fig 3A-B, Suppl. Fig 5A-B-C). Cells were differentiated and stimulated with blue light as previously described in Figure 2A and harvested at day 14 for single cell RNA-sequencing analysis. Differentially expressed genes based on leiden clustering, RNA-velocity trajectories and cell cycle predictions were used to classify 14 distinct cell identities: (i) FOXA2^+^/ARX Floor Plate; (ii) NKX2-1^+^/RAX^+^/SIX6^+^ Ventral tuberal hypothalamic progenitors; (iii) NKX2-1^+^/FOXG1^+^ Ventral telencephalic progenitors, (iv) NKX2-1^+^/NHLH2^+^/OTP^+^ Ventral hypothalamic neurons; (v) TFAP2A^+^/KRT19^+^ Superficial ectoderm; (vi) SOX10^+^/PLP1^+^ Neural Crest, (vii) PAX6^+^/EMX2^+^/OTX2^+^ Dorsal forebrain progenitors; (viii) IRX3^+^/ OLIG3^+^ Dorsal thalamic progenitors; (ix) TBR1^+^/LHX1^+^ Dorsal Neurons_1; (x) HES6^+^/DLL3^+^ Dorsal Neurons_2; dorsal and ventral proliferating progenitors, (xi) Dorsal, (xii) Dorsal thalamic, (xiii) Ventral and (xiv) an unidentified population (UnId) (Fig 3B, Suppl. Fig 5C-D, Supplementary Table 1). This analysis revealed the presence of multiple cell types that demarcate different domains along the embryonic D-V and A-P axes in agreement with what was previously shown by asymmetric SHH stimulation in 3D organoids (Cederquist et al., 2019). No endoderm, mesoderm or extra-embryonic markers were detected, locating cells in the ectodermal compartment.

To correlate the timing of our self-organizing *in-vitro* tissues with *in-vivo* events, we integrated our dataset with a scRNA-seq collection of mouse brain samples at different time points, E8.5, E10, E12, E12.5, E13 and E15 (La Manno et al., 2020). scRNA-seq transcriptomics of the human light-induced cells, grouped as neural precours (NPCs), floor plate, superficial ectoderm, neurons and UnId cells, integrate with the *in-vivo* mouse brain developmental atlas from La Manno et al., 2020 (Suppl. Fig 6A). The human NPCs display high correlation with the Radial glia population at E10, while the in-vitro derived neurons display high correlation with the mouse neuronal category at E12.5-E15 (Suppl. Fig 6B-C), suggesting a temporal match with human development at PCW4-5 (https://embryology.med.unsw.edu.au/embryology/index.php/Carnegie_Stage_Comparison).

The expression of GLI3, GAS1 and PTCH was used to identify SHH receiving cells. In agreement with literature, we identified a GLI3-GAS1^low^/PTCH1^high^ population as SHH stimulated while GLI3-GAS1^high^/PTCH^low^ cells as unstimulated (Fig 3C, Suppl. Fig 7A). We also validated the specific induction of SHH signaling in the NG-T2A-SHH line compared to a NG-CNTRL line by testing GLI1 and GLI3 expression at day 14 using qRT-PCR (Fig 3D). Among the SHH induced populations, scRNA-seq analysis confirmed the presence of a FOXA2^+^/ARX^+^, floor plate population and revealed the identity of four distinct NKX2-1^+^ populations (Fig 3B-C, E). The first is representative of the tuberal hypothalamic neural progenitors positive for NKX2-1, SIX6, SIX3, RAX (Fig 3C-E-F-G, Suppl. Fig 7C) (Shimogori et al., 2010; Martinez-Ferre and Martinez, 2012; Morales-Delgado et al., 2014), the second is the NKX2-1^+^ FOXG1^+^ population representative of the ventral telencephalic population (Fig 3E-F)(Maroof et al., 2013; Cederquist et al., 2019), the third is a ventral population of proliferating progenitors and the fourth is a small population of ventral neurons that we classified as hypothalamic neurons positive for NKX2-1, OTP, NHLH2 (Fig 3B-C, Suppl. Table 1) (Martinez-Ferre and Martinez, 2012; Morales-Delgado et al., 2014). Among the SHH unstimulated cells, we identified dorsal populations that consist of forebrain progenitors (PAX6^+^/EMX2^+^/OTX2^+^), thalamic progenitors (IRX3^+^/ OLIG3^+^), two neuronal populations (TBR1^+^/LHX1^+^ and HES6^+^/DLL3^+^), and non-neural ectoderm derivatives such as superficial ectoderm and neural crest (Fig. 3B, Suppl. Fig 5C, Suppl. Table 1). The non-neural ectoderm population derive mostly from the plate edge independently from the light organizer (Suppl. Fig 7B).

Immunostaining for specific markers, SIX6, RAX, NKX2-2 and FOXG1, revealed the spatial segregation of telencephalic and hypothalamic territory (Fig 3F-G, Suppl. Fig 7C-D-E). We further showed that light-modulation of SHH not only self-organize telencephalic and hypothalamic progenitors, but also neurons, since hypothalamic OTP^+^ neurons are preferentially located proximal to the light-induced organizer (Fig 3H). While the differentiation of hypothalamic cells has been previously observed in traditional cell culture or 3D organoids (Maroof et al., 2013; Merkle et al 2015; Qian et al., 2016; Cederquist et al., 2019; Kasai et al., 2020), confinement of a SHH source in 2D instructs the self-organization of a ventral telencephalic-hypothalamic structures that are spatially organized in monolayer.

Finally, the hypothalamic marker genes used in this study were *in-vivo* validated for their specific expression in the human fetal hypothalamus at PCW10 (Suppl. Fig 8A) (Zhou et al., 2020). In order to capture the gene expression modules that are shared between our *in-vitro* dataset and the fetal sample from Zhou et al., 2020, we compute the gene regulatory networks (regulons) in each dataset using pySCENIC (Van de Sande et al., 2020). Among the 427 active regulons identified in the human fetal hypothalamus dataset, the 72.5% (310 regulons) are shared with our *in-vitro* dataset (Suppl. Fig 8B, Suppl. Table 2). Based on RNA velocity analysis, we identified in the light-induced scRNA-seq dataset a ventral differentiation trajectory that starts from the ventral proliferating progenitors and ends at the ventral hypothalamic neurons (Fig 3B). Performing gene ontology analysis of genes that are differentially expressed along this trajectory, we identified waves of gene expression linked to cell-cycle regulation, neural progenitor expansion and neuronal maturation (Suppl. Fig 8C). We explored whether some important gene expression patterns recently described in the context of human fetal hypothalamic development were recapitulated in our model, TTYH1, HMGA2 and MYBL2 show the same progenitor-neuronal trend observed in Zhou et al., 2020 (Suppl. Fig 8C).

Collectively, our experiments demonstrate the ability to generate organizing centers by a simple blue light-induction of a morphogen, which specify a morphogenic source that pattern discreate cell types along a proximal distal axis in space, and lineage trajectory in time.

## Discussion

We devised a collection of optogenetic ePiggyBac vectors to conditionally express a photoactivatable Cre-loxP recombinase, that allows the control of morphogen expression using a simple blue light stimulation in hESCs. In this study, we used this tool to induce a localized domain of SHH expression to generate a ventral organizer within a neutralizing tissue. This establishes proof of feasibility for a new technological concept, which we predict to apply to a variety of gene expression modules, including signaling pathways, transcription factors and cell cycle players. This new experimental embryology tool has advantages over the classical techniques as it provides higher throughput and resolution. This system allowed us to control the asymmetric induction of the morphogen SHH during hESCs differentiation, which generated multiple locally organized cell types. It represents the first generation of light induced self-organization of cell fates and patterns in hESCs. Future optimization will improve several aspects. For example, it is foreseeable to add a pulsatile temporal regulation to this set up by imposing another layer of regulation rather than a step induction as it is in this study. Traditional methods for inducing local perturbation of signaling pathways, such as beads, provide little control over the extent of the inductive field’s and are incompatible with high-throughput setups. Our *in-vitro* model system displays high reproducibility and spatial control among many independent wells of differentiating human pluripotent stem cell. Since our light inducible experimental setup is unidirectionally activated, the forebrain D-V structures obtained in this study, are likely generated by the combination of direct SHH signaling and proliferation/expansion of the induced progenitors. We envision that, manipulating multiple signals using different light wavelength provides the promise of more sophisticated inductive interaction studies (Yu et al., 2020; Yen et al., 2020). For example, pairing ligands and inhibitors to precisely decipher the cellular and molecular aspects of a Turing based reaction diffusion events, which have been shown to have an instructing role in the establishment of embryonic patterns.

Light control of gene expression allows geometric confinement of the stimulus and its response in inducing the ventral organizer SHH. We have previously shown that physical geometrical confinement of hESCs is sufficient to induce self-organization in response to BMP4 in the context germ layers (gastruloids) (Warmflash et al., 2014). Self-organization has also been demonstrated to occur in the context of a single embryonic germ layer, ectoderm (Haremaki et al., 2019; Knight et al., 2019). Dual SMADs inhibition leads to self-organization of telencephalic neural progenitor around a lumen generating rosette (cerebroids) (Haremaki et al., 2019; Knight et al., 2019). When Dual-SMADs inhibition is followed by BMP4 presentation, confined hESCs colonies self-organize to generate the four derivatives of the ectodermal layers: neural, neural crest, sensory placode and epidermis that organize in radially symmetrical patterns (Haremaki et al., 2019). In the context of the ectoderm, this type of self-organization reflects the events that occur in the dorsal anterior part of the developing central nervous system. As for spatial geometrical confinement that provided the key element for gastruloid, cerebroid and neuruloids self-organization, here we confined the geometry of a morphogenic organizer by imposing a chemical edge with light stimulation. Complementing these results, our tool allowed the self-organization of the ventral aspect of the developing nervous system. It is tempting to speculate that self-organization imposed by geometrical confinement when combined with spatial manipulation of a morphogen will lead to more sophisticated and complex aspects of self-organization. Confinement of geometry and organizer structures promise to generate highly quantitative models that will more faithfully represent early human development *in-vitro*.

We and others have shown that it is possible to deconvolve several aspects of embryonic self-organization by modelling developmental events *in-vitro*. Our study uses a light inducible tool to generate 2D models of D-V forebrain development, composed of neural telencephalic and hypothalamic populations. Interestingly, the hypothalamus is among the most conserved structure of the brain in vertebrates. In the adult, it is composed of several spatially distinct nuclei that control a wide range of functions, from body homeostasis to behavior and it is connected with the endocrine system through the pituitary gland. To the best of our knowledge, a comprehensive and systematic induction of discreate hypothalamic nuclei in the human context has not been achieved yet. Spatiotemporal induction of specific signaling pathways in the context of hypothalamic self-organization promise to recapitulate, *in-vitro*, several aspects of the highly spatially organized hypothalamic development. Self-organization follows self-organization, beginning from an entire embryo (gastruloid) to a single embryonic germ layer such as the ectoderm (cerebroids and neuruloids), and ultimately the formation of discreate organs and cell types (dorsal and ventral brain). By this logic we expect that the development of discreate hypothalamic nuclei will also follow the rules of self-organization and symmetry breaking. We speculate that the generation and patterning of hypothalamic territories is possible as long as the appropriates physical (confinement) and chemical (morphogen) cues are provided. Self-organizing models of the human hypothalamus *in-vitro* will shed light on the basic principles that govern the development of a fundamental brain controlling center.

---

**Suppl. Fig 1:**
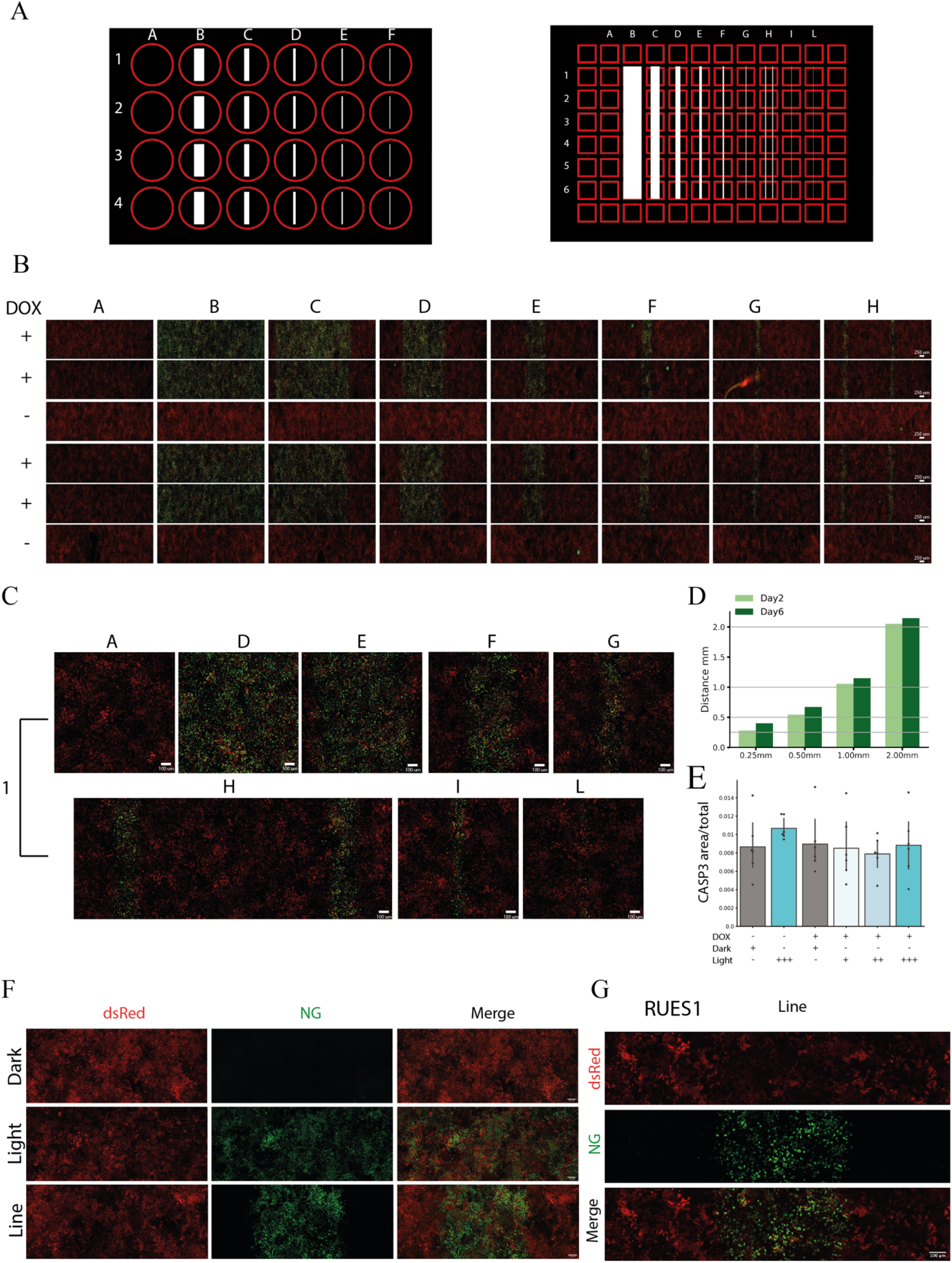
Systematic and robust light-induced gene expression in hESCs. **(A)** Blueprint of the film photomask used in combination with 24 (left) or 96 (right) multiwells plates with light-permeant aperture. Red label represents the well border while white label is the light permissive region. Light permissive regions for 24 well plate: A=0mm, B=4mm, C=2mm, D=1mm, E=0.5mm, F=0.25mm. Light permissive regions for 96 well plate: A=0mm, B=7mm, C=3.5mm, D=2mm, E=1mm, F=0.5mm, G=0.25mm, H=2x 0.25mm, I=0.125mm, L=0.625mm. **(B)** hESC treated with or without DOX and Blue-light for 24h hours, individual wells of a 96 well plate are displayed using the merge of Green (Green module) and Red (Red Module). Columns A to H and row 1 to 6, with light aperture: A=0mm, B=7mm, C=3.5mm, D=2mm, E=1mm, F=0.5mm, G=0.25mm, H= 2 lines of 0.25mm. **(C)** Confocal fluorescence images, Green (Green module) and Red (Red Module) show the use of light responsive hESC in combination with a 96-well plate set up for light patterning. Selected wells in columns A, D, E, F, G, H, I and L, from row number 1 display photomask patterning after DOX induction and 24h of Blue light. **(D)** Analysis of the Green module induced territory at different time points (day 2 and day 6) in relation with their corresponding photomask (N=2). **(E)** Quantification of the CASP3 positive area upon light-stimulation. Conditions: presence or absence of Dox with concomitant Dark or Light stimulation. We titrated the time of light-stimulation using different pulsed light conditions. 1 cycle of pulsed light is equal to 20 seconds of Light-ON and 120 seconds of Light-OFF. The light stimulation intervals are: 1 cycle, 25 cycles, and 600 cycles (n=5). **(F)** Confocal fluorescence images, Green module and Red Module of a 1mm Line induced area at day 6, under full Dark or Light. Red module (dsRed), Green module (NG) and merge composite channel. **(G)** Validation of optogenetic ePiggyBac vectors to conditionally express a photo-activatable Cre-loxP recombinase in an independent hESCs line: RUES1 (NIHhESC-09-0012). Confocal fluorescence images of photomask induced light patterns showing the expression of the Red module (dsRed), Green module (NG) and merge composite channels.

**Suppl. Fig 2:**
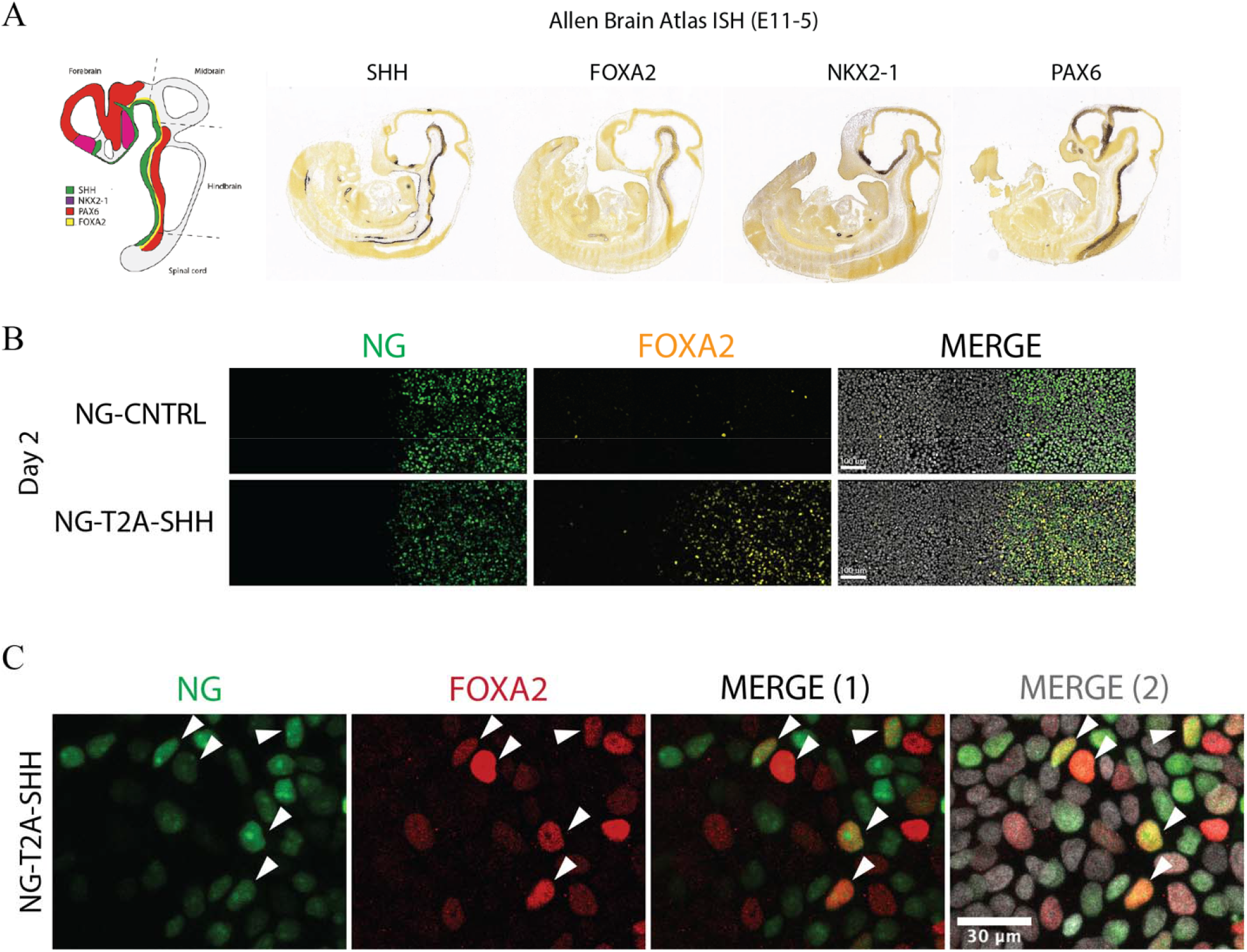
Ventral markers and autocrine induction of FOXA2 positive ventral cells upon light-induced SHH. **(A)** Schematization of mouse embryos in a parasagittal section for Dorso-Ventral (D-V) marker genes (Left panel). Allen brain atlas ISH showing D-V marker genes in the mouse embryo E11.5 (https://developingmouse.brain-map.org) (NKX2-1, SHH, PAX6, FOXA2). **(B)** Fluorescence images of large fields of view showing consistent FOXA2 induction along the light-induced line of SHH, but not the control NG-CNTRL at day 2. Color code is: NG-Green (Organizer), FOXA2-Yellow, DAPI-Grey and Merge-Composite. Scale bar: 100μm. **(C)** Immunostaining at higher magnification showing double positive NG-SHH and FOXA2 cells. DAPI-Grey, NG-Green, FOXA2-Red and Merge-Composite. Scale bar= 30μm.

**Suppl. Fig 3:**
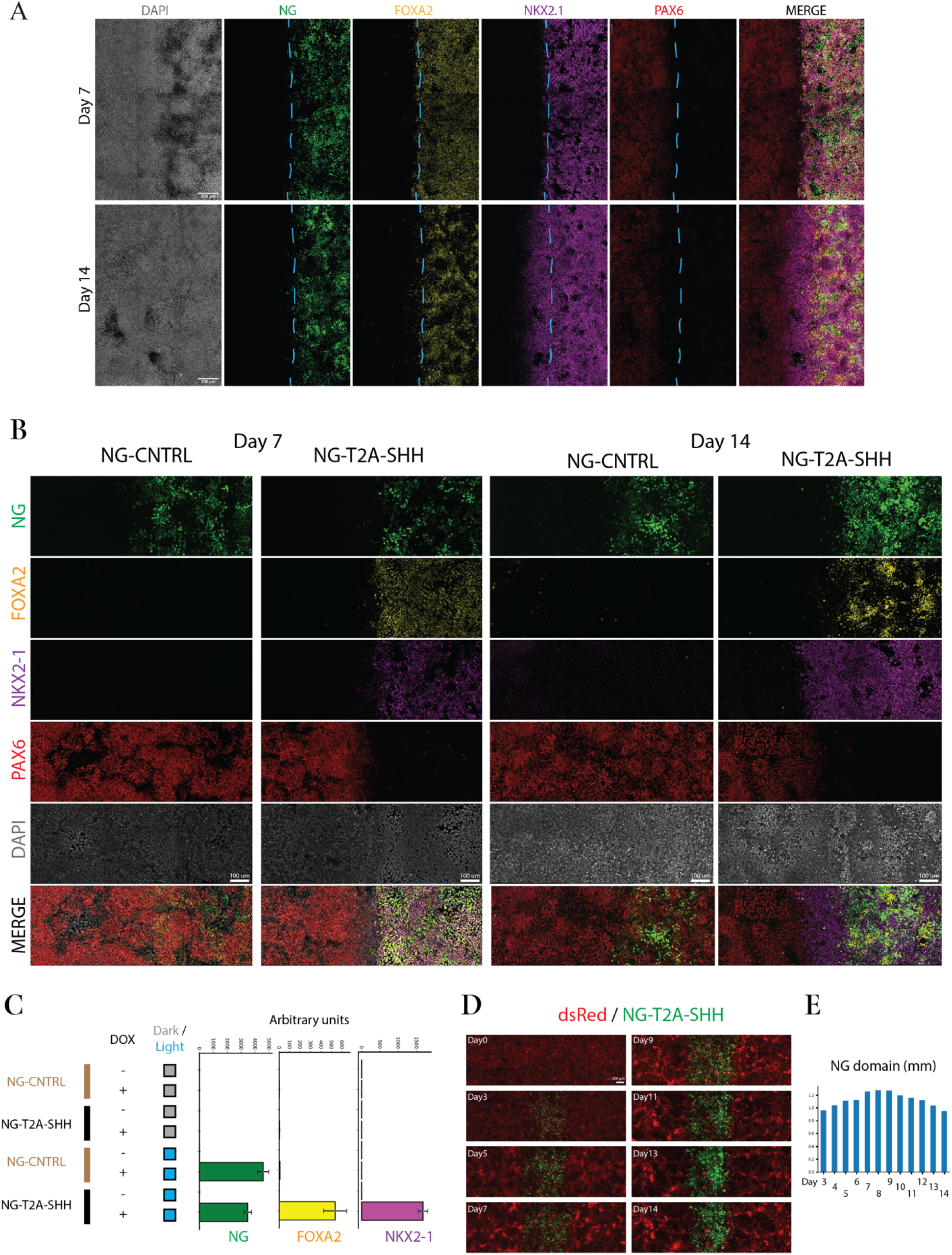
SHH-dependent dorsal-ventral self-organization. **(A)** Immunofluorescence images of large fields of cells showing consistent patterning along the light-induced line of SHH at day 7 and day 14. Color code is: NG-Green (Organizer), FOXA2-Yellow (Ventral), NKX2-1-Magenta (Ventral), PAX6-Red (Dorsal), Merge-Composite. Dashed cyan line indicate the border of SHH producing cells. Scale bar: 200μm. **(B)** Immunostaining for dorsal-ventral markers in light induced SHH cells and NG-CNTRL during differentiation at different time points, day 7 and 14. DAPI-Grey, NG-Green, FOXA2-Yellow, NKX2-1-Magenta, PAX6-Red and Merge-Composite. Scale bar: 100μm. **(C)** qRT-PCR showing the mRNA level of ventral marker genes (FOXA2 and NKX2-1) specifically induced in the NG-T2A-SHH line but not in the control line, NG-CNTRL. Dark condition and DOX-minus controls do not induce ventral marker genes during differentiation. Histograms display the average and the standard deviation (n=2). D) Time-laps imaging of NG-T2A-SHH cells induced with a 1mm photomask. Images display day: 0,3,5,7,9,11,13,14. dsRed-Red and NG-green. Scale bar 200 μm. E) Analysis of the Green module induced territory at different time points (day: 3,4,5,6,7,8,9,10,11,12,13,14)

**Suppl. Fig 4:**
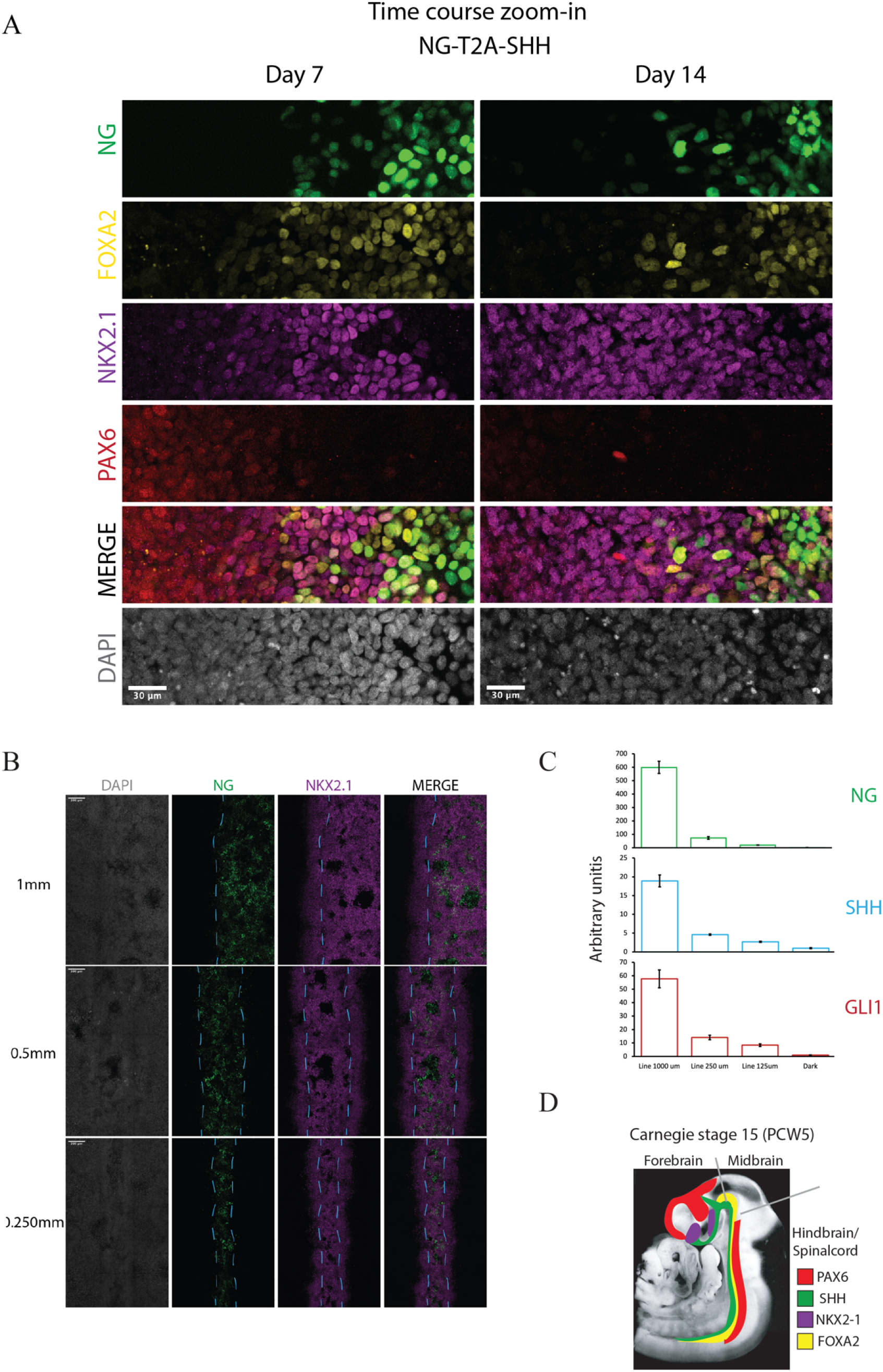
Stimulus-dependent expression of SHH targets upon light induction. **(A)** Immunostaining at higher magnification showing details of ventral fates induced in proximity of the NG-T2A-SHH organizer at day 7 and 14. DAPI-Grey, NG-Green, FOXA2-Yellow, NKX2-1-Magenta, PAX6-Red and Merge-Composite. Scale bar: 30μm. **(B)** Fluorescence images of large fields of view showing consistent patterning along the light-induced line of SHH at day 14. Color code is: NG-T2A-SHH-Green (Organizer), NKX2-1-Magenta (Ventral) and Merge-Composite. Dashed cyan line indicate the border of SHH producing cells. Scale bar: 200μm. **(C)** qRT-PCR showing the mRNA levels of NG (Green module), SHH, and the SHH-responsive gene GLI1, in samples spatially activated with light using a photo-mask of: 1000μm, 250μm and 125μm. NG, SHH, and its downstream target correlate with the area of induction. Histograms display average and standard deviation (n=2). D) Carnegie collection image featured with a color-coded scheme of the human sagittal section Carnegie stage 15 (PCW5) for specific markers (SHH-Green, NKX2-1-Magenta, PAX6-Red, FOXA2-Yellow), dashed lines delineate Forebrain, Midbrain, and Hindbrain / Spinal cord boundaries.

**Suppl. Fig 5:**
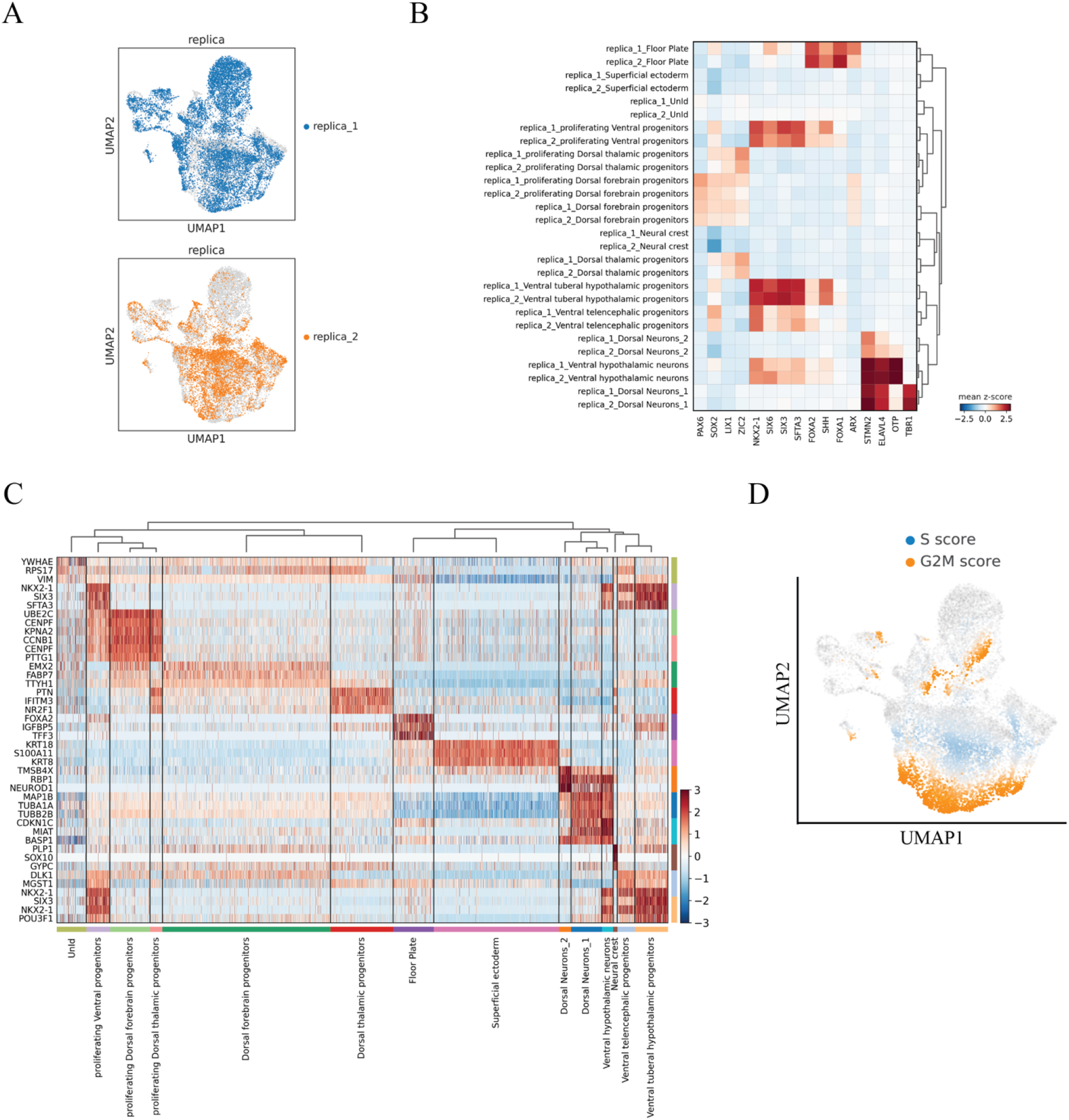
scRNA-seq transcriptomic analysis of SHH-induced fates. **(A)** UMAP plot of the two-biological replica of *in-vitro* light induced SHH cells at day 14. The two replicas integrate in the UMAP space and display the same cellular populations in both replicas. **(B)** z-score heatmap of the two replicas showing the consistent expression in the independent replicas of marker genes used for classification. **(C)** z-score heatmap display the top 5 marker genes for each cell identity in the light induced SHH scRNA-seq dataset. **(D)** scVelo calculated cell cycle prediction were embedded in the UMAP space and used for classification of the proliferating cells.

**Suppl. Fig 6:**
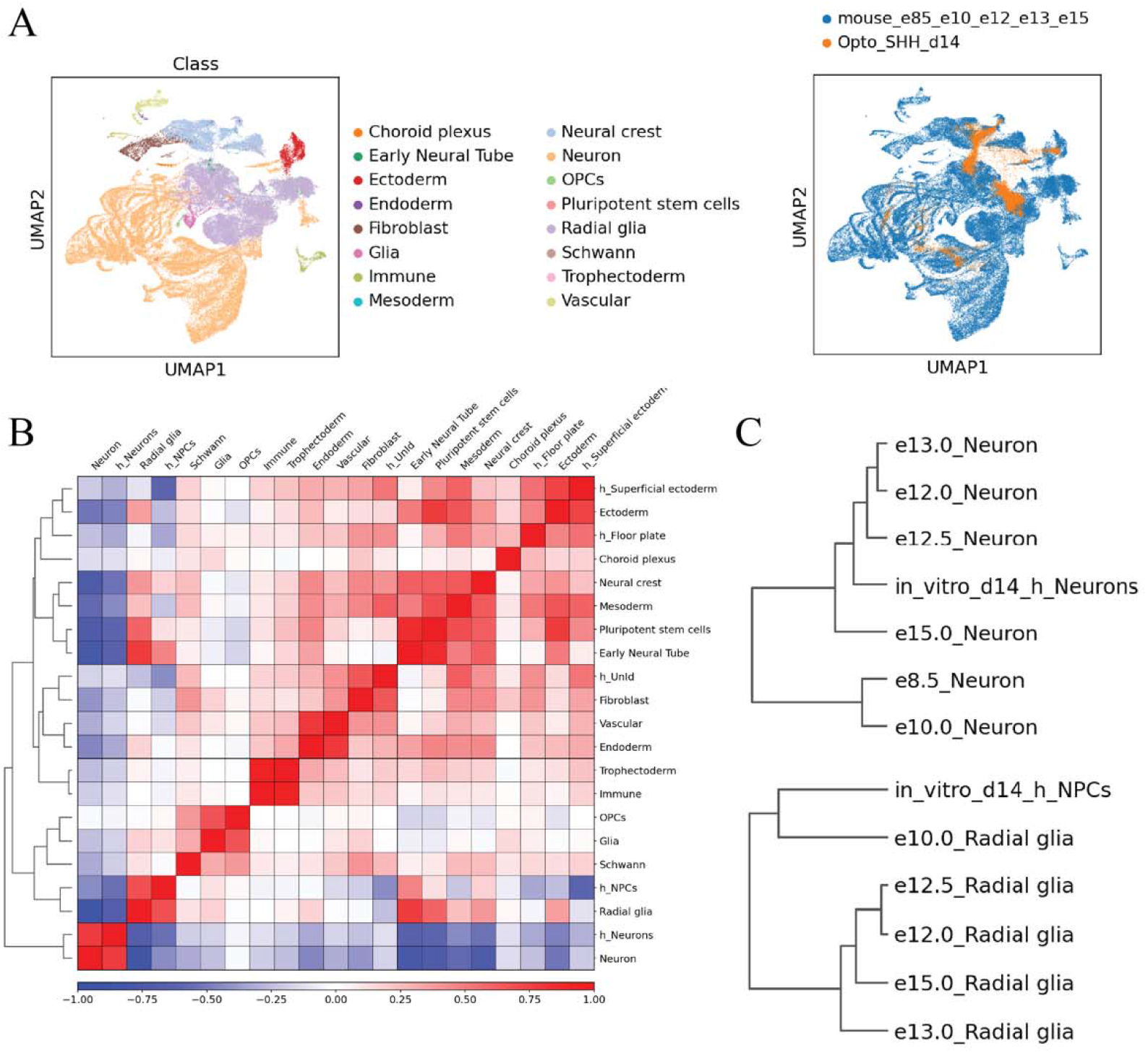
Conserved gene expression between light-induced synthetic tissue and their mouse embryonic counterparts. **(A)** UMAP plot of the mouse brain dataset from La Manno et al., 2020 showing cell populations at different time points (E8.5, E10, E12, E12.5, E13, and E15), UMAP plot shows the mouse dataset annotation “Class” (top left). UMAP plot of the human *in-vitro* light induced SHH cells integrated with the mouse dataset in defined clusters (top right). The human cells mostly integrate with the E10 time points in the Radial glia population, the Ectoderm, the Neural Crest and Neuronal label annotations. (B) Correlation analysis using the high variable genes of the human-mouse integrated dataset displayed as z-score heatmap using the mouse “Class” annotation and the human light induced cells, grouped as neural precursor (NPCs), floor plate, neurons and UnId cells. **(C)** Dendrogram analysis shows clustering of the *in-vitro* derived human cells grouped as NPCs and Neurons, with specific time points of mouse brain development.

**Suppl. Fig 7:**
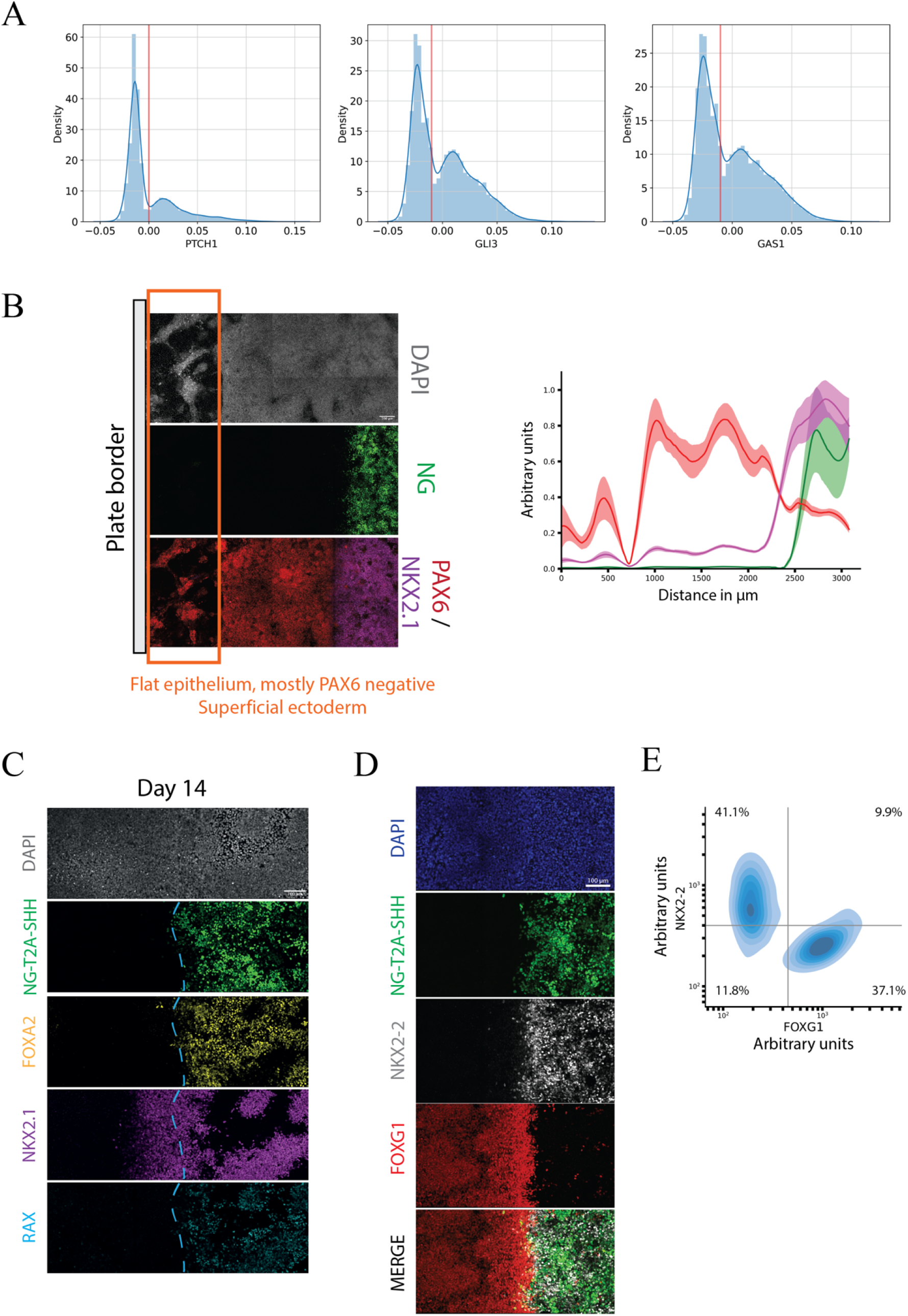
Identification of SHH responsive cells. **(A)** Distribution of log normalized scRNA-seq counts for selected genes: (PTCH1, GLI3 and GAS1). Based on the bimodal distribution, specific thresholds are manually selected and displayed as Red vertical bar. Thresholds are used to categorically labels cells as SHH stimulated when GLI3-GAS1^low^/PTCH1^high^ conditions are valid. **(B)** Large field of view display the well edge that generate most of the PAX-negative cells which correspond to superficial ectoderm and neural crest independently from the light induced SHH organizer (left panel). Cumulative fluorescence intensity analysis (line profile) over the x-axis from the edge of the 96 well. x-axis displays the linear distance in μm. y-axis shows the cumulative fluorescent intensity profile in arbitrary units for each channel. Line profile shows the average (line) and standard deviation (area) for each channel. The line profile is color-coded as the immunofluorescent channels, NG-Green, NKX2-1-Magenta, PAX6-Red at day 14 quantification (n=4) (right panel). **(C)** Immunostaining shows the spatial segregation of ventral population arising from a light induced SHH source using an additional hypothalamic marker RAX at day 14. (Green NG, Yellow FOXA2, Magenta NKX2.1, Cyan RAX, Grey DAPI), Scale bar = 100μm. D) Immunostaining shows the spatial segregation of telencephalic (FOXG1) and hypothalamic (NKX2-2) marker in relation to the light induced SHH source of SHH at day 14. (Green NG, Grey NKX2-2, Red FOXG1, Blue DAPI), Scale bar = 100μm. E) Immunostaining single nuclei quantification of NG-T2A-SHH induced cells displayed as density plot at day 14. x-axis report FOXG1 while the y-axis the NKX2-2 fluorescence intensity profile using arbitrary units in a log_10_ scale.

**Suppl. Fig 8:**
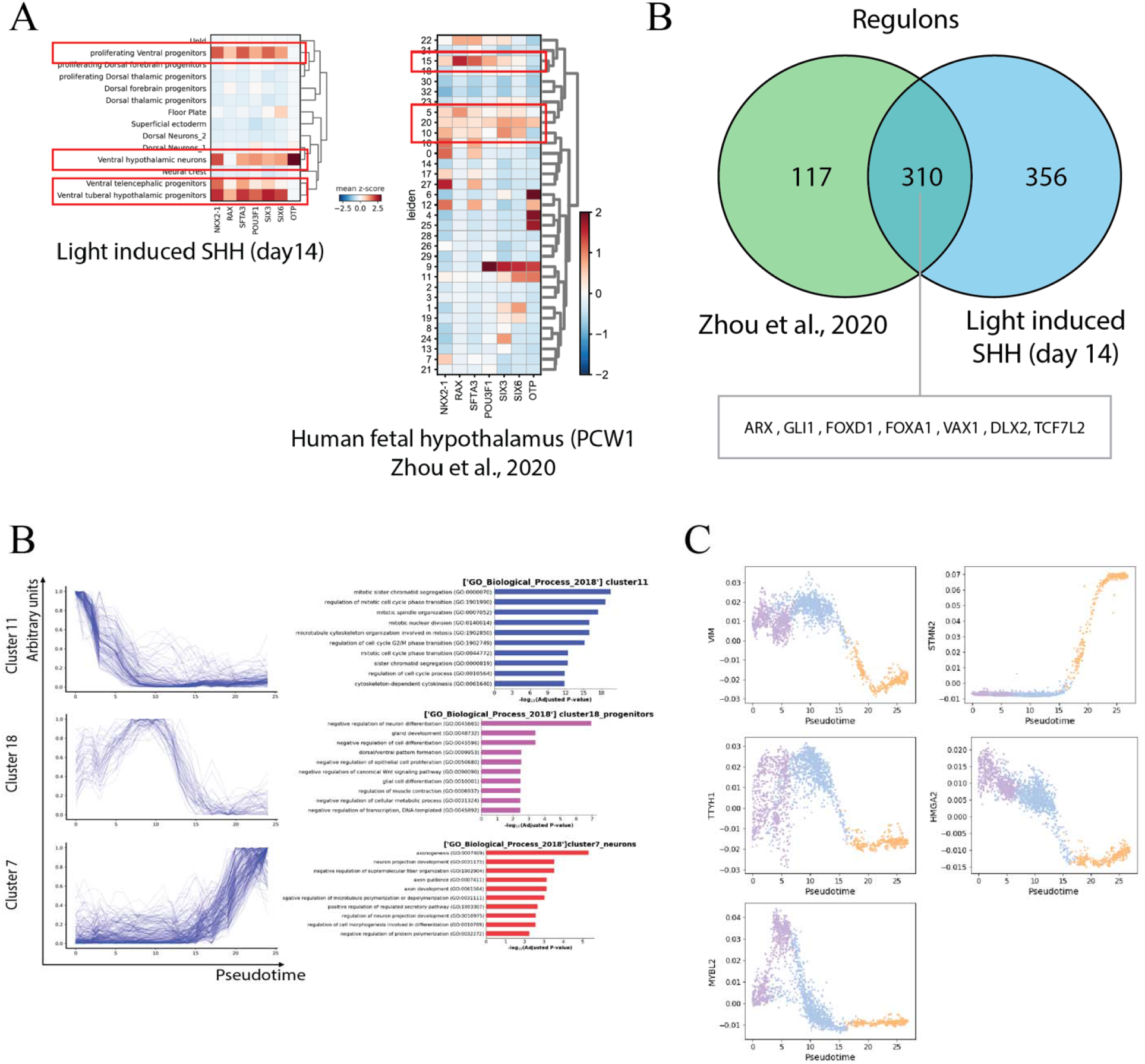
Conserved gene expression between light-induced synthetic tissue and their human fetal hypothalamic counterparts. **(A)** Comparative analysis of hypothalamic markers detected in our light-induced SHH scRNA-seq data, with that of hypothalamic human fetal samples (PCW10; Zhou et al., 2020). **(B)** Venn-diagram of the active regulons commonly identified using pySCENIC (Van de Sande et al., 2020) between the fetal hypothalamic dataset (Zhou et al., 2020) and the light induced SHH dataset at day 14. **(C)** Selection of gene patterns identified along the hypothalamic ventral trajectory displayed as line plot over pseudotime, each line represents a gene. The x-axis corresponds to the pseudotime time along the trajectory and the y-axis to the normalized 0-1 scaled gene expression values. Gene patterns have been tested for enrichment of specific categories using GSEAPY-enrichr (GO biological process 2018). **(D)** Pseudotime analysis in the light induced hypothalamic ventral trajectory for selected candidates: VIM, STMN2 are markers used to validate the progenitor to neuron trajectory (Suppl.Fig 8, top panel); TTYH1,HMGA2 and MYBL2 are candidates described in the fetal hypothalamic development (Zhou et al., 2020). TTYH1, HMGA2 and MYBL2 are downregulated along the differentiation trajectory in our *in-vitro* model, in agreement to what was recently shown in the human fetal hypothalamus (Zhou et al., 2020). The scatterplot displays individual cells from the subset of NKX2-1^+^ cells in light induced scRNA-seq dataset. The x-axis reports the pseudotime time along the ventral trajectory and the y-axis the normalized expression values.

## Material and methods

### Human pluripotent stem cell culture

hESCs lines used in this study have been described previously in James et al., 2006, and are part of the NIH Human Embryonic Stem Cell Registry (RUES1-NIHhESC-09-0012) and (RUES2-NIHhESC-09-0013). Human pluripotent cells were maintained in feeder-free conditions on Geltrex-coated dishes. Cells were feeded with MEF conditioned medium (CM) supplemented with bFGF 20ng/mL daily (James et al., 2006). Cells were routinely passaged every 3-4 days using Gentle Cell Dissociation Reagent (STEMCELL-Technologies).

### Neural induction for pluripotent stem cell

hESC were passaged as single cells using Accutase (STEMCELL-Technologies) and seeded in CM media supplemented with bFGF and Y27632 at 100,000 cells/cm^2^ density. The differentiation started 1-2 days later when cells reached 100% confluency. Neural induction media is composed of: 50% Neurobasal and 50% DMEM-F12, supplemented with N2 0.5%, B27 0.5%, Glutamax 0.5X and Non-essential amminoacid 0.5X and Insulin (2.5μg/ml) all from LifeTechnologies. Small molecule SB431542 (10μM) and LDN193189 (100nM) were supplemented to warm media.

DOX (1μg/ml) induction is used at day “0” concomitantly with dual-SMAD inhibition, and then washed out at day “2”. Neural induction media supplied with dual-SMAD inhibitors is maintained throughout differentiation until day “14”.

### piggyBac vectors generation

One of the two piggyBac DOX inducible and Puromycin selectable vector (Rosa et al., 2014) was digested with HindIII-NotI and ligated with the Magnets-CRE expression cassette from Kawano et al., 2016 (pcDNA-MagnetsCRE plasmid is a kind gift from Dr. Yazawa). The other that is Blasticidine selectable, and carries a constitutive promoter (CAG), was modified to express a LoxP regulated switch of the two ORFs. A first “Red module” that express a dsRed protein, is removed and upon LoxP recombination allows the expression of a second “Green module”, which express a Neon-Green protein fused with a T2A, allowing co-expression with other proteins (Fig 1B). The LoxP-dsRed-LoxP was cloned using BamHI-BglII restriction site from the pLV-CMV-LoxP-DsRed-LoxP-eGFP, gift from Jacco van Rheenen (Addgene plasmid # 65726). PCR amplified SV40 poly-adenylation site was introduced using NheI restriction enzyme after the dsRed ORF. The Green module consisting of the nuclear localized Neon-green protein was synthesize using IDT geneblocks and inserted using the restriction enzymes AgeI-NotI. The SHH coding sequence was PCR amplified and cloned using BsmBI-NotI restriction enzymes from pOEM1 pCMV:hShh-pPH:vsvged was a gift from Elly Tanaka (Addgene plasmid # 111156). Plasmids used in this study have been deposited on ADDGENE.

### Stable integration of piggyBac vectors in the genome of hESCs

hESCs were nucleofected using an Amaxa Nucleofector II (Lonza) according to manufacture instruction for hESC nucleofection. A mixture (1μg+1μg+0.5μg) of: ePB-PURO-TT-PA-CRE, ePB-BSD-CAG-RLOXP-NG-T2A or ePB-BSD-CAG-RLOXP-NG-T2A-SHH, and the piggyBac transposase were nucleofected in 10^6^ hESCs and plated in Geltrex (Life technologies) coated plates using CM media supplemented FGF2 (20ng/mL) and ROCK-inhibitor (Y-27632, 10 μM) for 24 hours. Cells were then selected using CM media supplemented with blasticidin 5μg/ml and puromycin 1ug/ml (all from Life technologies) for 2 weeks.

### Immunostaining

Cells were washed in PBS-/- and fixed in PFA 4% for 30 minutes at room temperature (RT). After fixation, 96 or 24 wells plates have been washed three times with PBS-/-. Blocking was performed in PBS-/-, TritonX (0.15%), Normal Donkey Serum 3% (Ab-solution) for 30 minutes at room temperature. Primary and secondary antibodies staining was done at room temperature for 1 hour each, in Ab-solution, followed by 3 washes in PBS-/- TritonX (0.15%). The list of primary antibodies is provided below. DAPI (Thermo Fisher Scientific) staining was performed 5 minutes and plates were stored in IBIDI mounting medium at 4 degree Celsius before imaging.

### Blue light-stimulation

Cells were plated into IBIDI 24 or 96 wells plates and placed on top of a custom-made black plastic box. The bottom on the IBIDI plate was aligned with a custom-made laser cut plastic layer or with a film photomask that spatially define the area of illumination. The Blue light sources used in this study are a custom LED-blue light tablet or the LED 96 well array system (AMUZA INC). Pulsed illumination was delivered to the samples with the following parameters: 20sec ON-time, 120sec OFF-time cycling for 24 hours at 10V power.

### scRNA-seq preparation

scRNA-seq sample preparation was performed following manufacture instructions. In brief, cells were enzymatically dissociated as single cells using Accutase for 8 minutes at 37°C, washed twice in PBS-/- BSA 0,04% and then strained with a 40μm filter. Gem formation and library preparation was performed following manufacture instructions using Single Cell 3′ v.3 reagents (10X Genomics).

### scRNA-seq analysis

The scRNA-library for light induced SHH at day 14 during neural induction was sequenced using a NovaSeq 6000 SP flowcell as 28 × 94 × 8. FASTQ file were aligned using Cell Ranger (v.2.0.2) against hg19 reference genome. Count matrix were further processed in Python using Scanpy environment (Wolf et al., 2018)(https://pypi.org/project/scanpy/) and are available under the NCBI GEO accession number GSE163505. The raw matrices were filtered to have a minimum of 500 detected genes per cell and genes were filtered to be expressed in at least 5 cells. Cells with over 15% mitochondrial UMIs were discarded. Clustering analysis was performed using Leiden algorithm on normalized counts and visualized in a low-dimensional space using UMAP plots (n.neighbors□=□50, PCA components=20). Marker genes used to annotate the clusters were identified using a Wilcoxon rank-sum (Mann-Whitney-U). The integration of scRNA-seq biological replicas has been performed by matching mutual nearest neighbors using the Scanpy external API MNN correct, with default parameters (svd_dim=50) (Haghverdi et al., 2018).

### RNA velocity and Pseudotime analysis

Kallisto Bustools (Melsted et al., 2021) was used to align and quantify layered spliced and unspliced gene counts. The two scRNA-seq replicas were aligned independently, using kb-python=0.24.4 to capture spliced and unspliced transcripts, applying the --lamanno function with the GRCm38 human genome. The Kallisto unfiltered count matrix was imported in Scanpy and further processed using scVelo (v0.2.2) (Bergen et al., 2020), using UMAP coordinates and clusters annotation from previously annotated Scanpy object. Count matrixes carrying the spliced/unspliced layer annotation were filtered and normalized for minimum shared counts=30 and the top 2000 high variable genes. The first 30 principal components and 30 neighbors were used to calculate ‘Ms’ and ‘Mu’ moments of spliced/unsliced abundance. We used the steady-state model (from La manno et al., 2018) to determine velocities using the “scv.tl.velocity” function with default parameters.

We computed pseudotime analysis on a subset of cell classified as NKX2-1^+^ hypothalamic cells, consisting of: NKX2-1^+^/RAX^+^/SIX6^+^ Ventral tuberal hypothalamic progenitors, NKX2-1^+^/NHLH2^+^/OTP^+^ Ventral hypothalamic neurons and NKX2-1^+^/TOP2A^+^ ventral proliferating progenitors. Using the python wrapper of Slingshot, “scprep.run.Slingshot” (https://pypi.org/project/scprep/), we calculated the pseudotime order of cells using full covariance matrix and default parameters. The normalized and log transformed count matrix for gene trends investigation has been denoised using MAGIC (van Dijk et al., 2018) (default parameters, t=3). Trajectory scatterplot are displayed in function of the pseudotime, x-axis, and the gene expression values, y-axis.

Gene patterns have been identified using the scaled high variable genes and the Agglomerative Clustering algorithm (sklearn, https://scikit-learn.org/stable/) along the pseudotime (clusters number=20) using default parameters. Selected gene pattern clusters have been tested for enrichment of specific categories using GSEAPY-enrichr (GO biological process 2018)(https://pypi.org/project/gseapy/, Chen et al., 2013).

### Analysis and integration of publicly available scRNA-seq datasets

scRNA-seq samples from the mouse brain developmental atlas (La manno et al., 2020) were downloaded as Loom file and sliced according to their column annotation (E8.5, E10, E12, E13, E15). Extracted count matrixes were further normalized, log transformed and filtered for mitochondria content less than 15% and genes to be expressed in at least 5 cells. Gene annotations were converted to their human orthologous using “scanpy.queries.biomart_annotations”. UMAP coordinates were calculated using (n.neighbors□=□50, PCA components= 30). Human and mouse datasets were integrated using Ingest “sc.tl.ingest” function, using the common expressed high variable genes between the two datasets, using default settings. Correlation analysis of the integrated dataset was performed using the “Class” annotation from the mouse annotation and the human light induced cells, grouped as neural precours (NPCs), floor plate, superficial ectoderm, neurons and UnId cells. Correlation analysis is calculated using the scanpy correlation matrix function with default pearson parameters. Dendrogam is calculated using default settings and 30 PCA.

scRNA-seq count matrix of the human fetal hypothalamus (Zhou et al., 2020) were preprocessed using the following settings: genes expressed in more than 5 cells and cells with less than 20% of mitochondrial genes were used for further analysis. Count matrixes have been normalized and log transformed. UMAP coordinates and clusters were obtained using (n.neighbors□=□10, PCA components= 30) and leiden clustering (resolution=1).

### Regulons identification

pyScenic is used to identify gene regulatory network in scRNA-seq dataset, using default parameters (Van de Sande et al., 2020). In brief, count matrixes were preprocessed using the following parameters: minimum number of genes per cell = 200 and genes expressed in at least 3 cells. The filtered count matrix is used to infer co-expression modules using the human transcription factors ranked database (https://resources.aertslab.org/cistarget/databases/). Arboreto algoritm (grnboost2) is used to calculate adjacence matrixes. Regulons are computed using the adjacencies matrixes of each dataset independently, followed by the prune module for targets with cis regulatory footprints (RcisTarget). Commonly detected regulons between the fetal hypothalamic dataset (Zhou et al., 2020) and our scRNA-seq dataset have been tested for enrichment of gene ontology, using the “Tissues protein expression from the human proteome map” from GSEApy-enrichr.

### Imaging

Confocal images were acquired on a Zeiss Inverted LSM 780 laser scanning confocal microscope with a ×20 dry objective.

### Imaging analysis

Images were preprocessed to display the Maximum Intensity Projection (MIP) of at least 4 z-stacks using ZEN black software. MIP images were subsequently imported into Python using the Czifile package. Line profile analysis consists of a single channel exported and the intensity values for each pixel over the y-axis were summed and displayed as a curve. Relative internal normalization is performed when indicated. Line profile in a 0-1 range was obtained normalizing values for each channel according to their maximum and minimum value.

Single cell fluorescence intensity quantification was performed by identification of individual cells using an Otsu binarized DAPI image followed by distance transform and Watershed algorithm (scikit-image) to separate overlapping nuclei. Pixel intensity of each object is plotted using the Seaborn python library.

Cell death is quantified by calculating the area of CASP3 positive staining normalized on the total area. CASP3 positive signal is obtained upon Otsu thresholding of CASP3 staining using the python package (scikit-image, threshold_otsu).

### qRT-PCR

RNA from individual wells was extracted using the Qiagen RNeasy Plus Mini Kit. 1ug of RNA was retrotranscribed using Transcriptor First Strand cDNA Synthesis Kit (Roche 04897030001) and Real-time quantification was performed using SYBR Green Master Mix (Roche 04887352001). HPRT or ATP5O endogenous control is used for internal normalization and results are expressed as fold change over a reference sample.

### Sanger sequencing

PCR-amplified amplicons (Hotstart Q5 polymerase, NEB) were sanger sequenced using Genewiz sanger sequence service under standard manufacture instructions.

#### Primers sequence

HPRT_FW: GCCCTGGCGTCGTGATTAGT

HPRT_RV: GGCCTCCCATCTCCTTCATCA

MAGNETS_FW: CTCTGACAGATGCCAGGACA

MAGNETS_RV: GATCAGCATTCTCCCACCAT

CAG_FW: GCTTCGAATTCTGCAGTCGACCG

NEON_GREEN_RV: GCTGGGAGAGAGGCCATGTTATCC

GLI1_FW: TGGCATCCGACAGAGGTGAG

GLI1_RV: CAGTTATGGGCCAGCCAGAGA

GLI3_FW: TTGCACAAAGGCCTACTCGAGACT

GLI3_RV: CTTGTTGCAACCTTCGTGCTCACA

SHH_FW: AATGTGGCCGAGAAGACCC

SHH_RV: AGATGGCCAAAGCGTTCAAC

ATP5O_FW: ACTCGGGTTTGACCTACAGC

ATP5O_RV: GGTACTGAAGCATCGCACCT

FOXA2_FW: CCGTTCTCCATCAACAACCT

FOXA2_RV: GGGGTAGTGCATCACCTGTT

NKX2-1_FW: AGGGCGGGGCACAGATTGGA

NKX2-1_RV: GCTGGCAGAGTGTGCCCAGA

### Antibodies list

Anti-PAX6 mouse monoclonal, BD Biosciences 561462, 1:100

Anti-TTF1 antibody [EP1584Y], Abcam, (ab76013), 1:500

Anti-HNF-3BETA/FOXA2, Neuromics, GT15186, 1:200

Anti-NKX2-2, DSHB, 74.5A5, 1:200

Anti-FOXG1 [EPR18987], Abcam, ab196868, 1:100

Anti-OTP [EPR22178-17], Abcam, ab254267, 1:200

Anti-SIX6, Abcam, ab251658, 1:200

Anti-Rx, Takara, M229, 1:200

## Author contribution

R.D.S. conceived the project, performed experiments and analyzed the data. E.F. contributed to experimental discussions, imaging analysis and to establish the LED-light setup. E.A.R. contributed to scRNA-seq analysis. A.H.B. supervised the project and secured funding. R.D.S and A.H.B. wrote the manuscript with input from all authors.

## Acknowledgments

We would like to thank Zeeshan M. Ozair for discussion, criticisms, advises and comments on the manuscript. Manon Valet for computational suggestions and Sandra Moreu for supporting plasmids generation. We also would like to thank Jean-Marx Santel and Peter Ingrassia for revising the manuscript. We are grateful to Brivanlou and Siggia lab members for inputs and criticism. We also would like to thank the Genomics Resource Center and the Bio-Imaging Resource Center for technical support. We wish to thank Kavli-foundation for supporting this research with the Kavli-NSI-Pilot grant to RDS and AHB. RDS was supported by EMBO-LTF-254-2019.

